# Second harmonic generation polarization microscopy as a tool for protein structure analysis

**DOI:** 10.1101/338137

**Authors:** Junichi Kaneshiro, Yasushi Okada, Tomohiro Shima, Mika Tsujii, Katsumi Imada, Taro Ichimura, Tomonobu M. Watanabe

## Abstract

Second-harmonic generation (SHG) is a nonlinear coherent scattering process that is sensitive to molecular structures in illuminated materials. We report SHG polarization measurement for the detection of protein conformational changes in solutions of macromolecular protein assemblies such as microtubules and protein crystals. The results illustrate the potential of this method for protein structural analysis in physiological solutions at room temperature without labelling.

---

Second-harmonic generation (SHG) microscopy is a powerful modality that enables the visualization of fibre structures without labelling. Such structures include the fibrillar collagen in tendons, myosin thick filaments in myosarcoma, and the densely bundled microtubules in living specimens^1-3^. SHG is a coherent process in which two photons that have the same energy are effectively ‘combined’ to generate a new photon with twice the energy; it was first discovered in 1961^4^. Twenty-five years later, SHG was used to determine the orientation of collagen fibres in a rat tail tendon^5^. Because SHG is sensitive to static electric field, SHG microscopy was also used to measure the action potential of a nerve cell^6,7^. The microscope design is based on that used for laser-scanning confocal fluorescence microscopes, so that the simultaneous observation of fluorescence and SHG is possible^8^. SHG is now also used for whole-animal imaging *in vivo*^9,10^.

The nonlinear susceptibility of a material to SHG is expressed as the 3^rd^-rank tensor (SHG tensor) originating from molecules with broken centrosymmetry and their alignments in macromolecular assemblies^1,2^. This unique feature may enable the acquisition of data about the spatial distributions of filament orientations, and can provide information on protein secondary structures, such as the axial pitch of alpha-helices^11^. Furthermore, SHG microscopy can be used to visualize the spatial distribution of fibre polarity directions in tissues with interferometric configurations^12,13^. Therefore, the SHG signal potentially carries structural information about proteins, which can be recorded optically without labelling.

For many decades, the major tools of protein structural analysis have been cryo-electron microscopy (cryo-EM) and X-ray crystallography^14,15^, both of which are currently advancing to the next generation^16,17^. Whereas cryo-EM and X-ray crystallography provide static information on protein structure, structural dynamics can be investigated using nuclear magnetic resonance (NMR) and Förster resonance energy transfer (FRET)^18,19^. These technologies seem to provide a comprehensive tool-kit for the analysis of protein structures. However, these tools present several important practical difficulties: biological samples should be preserved in a frozen-hydrated state prior to examination using an electron microscopy grid in cryo-EM observations; X-ray crystallography requires a highly-purified single crystal with a typical size of 30–300 μm; and NMR and FRET require chemical labelling. In contrast, SHG observation requires only a small (200–500 nm) crystal or fabric structure. In the present study, we propose bypassing the problems mentioned above by adding the measurement of SHG polarization to the tool-kit of structural analysis methodologies.

## RESULTS

### Improvement of signal-to-noise ratio in SHG detection

A microtubule (MT), which is a helix filament composed of heterodimers of α- and β-tubulin^20,21^, provides the best model for proof-of-concept because the structural differences between the nucleotide-dependent states in MTs have been studied extensively^22-26^. In order to obtain the SHG polarization dependence curve for the structural difference in the MT, we constructed a transmission-type SHG microscope with a rapid polarization control device comprising a pair of electro-optic modulators in the incident light pathway^27^ (Fig 1). In an observation of fixed nerve cells, we selectively observed the bundled MTs in the dendrites, and confirmed the electric polarity of the centrosomes using the present SHG microscope (Fig. S1).

**Figure 1.**
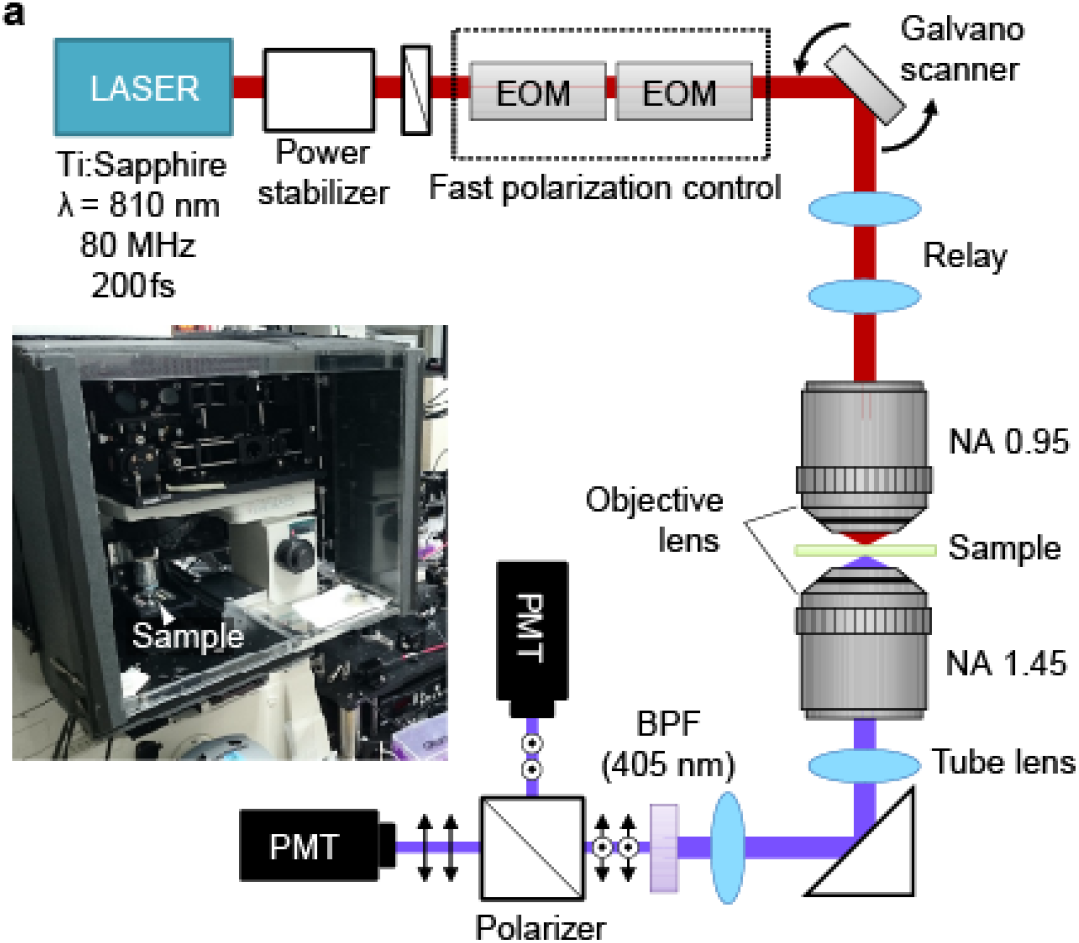
Configuration of our SHG microscope. Schematic drawing of the present SHG microscope. EOM, electrooptic modulator; PMT, photomultiplier tube; BPF, bandpass filter; NA, numerical aperture. The inserted photo is the present microscope combined with an incubator to stabilize MT polymerization.

A single MT fibre was preferred to simplify the interpretation of the experimental data. Although the sensitivity of the SHG detection of the present microscope was high enough to detect the SHG scattered from a single MT, the background noise degraded the signal-to-noise (S/N) ratio of the obtained image (Fig. 2a, *left*). SHG light generated in the glass coverslip on which the MTs were adsorbed was a major source of noise. Because the SHG from a commonly used borosilicate glass (known as BK7) exhibited non-negligible intensity as strong as that from a single MT (Fig. 2b, *left* and Fig. 2c, *left*), we replaced the borosilicate glass with high-purity SiO_2_ glass, which suppressed the SHG noise from the substrate (Fig. 2b, *right*). This improved the S/N ratio by approximately 2.8-fold (Fig. 2c), enabling the detection of the SHG signal from individual MTs (Fig. 2a, *right*). Therefore, the present method provided clear SHG images of individual MTs, even in fixed cells, and was at least as effective as fluorescence microscopy (Fig S2).

**Figure 2.**
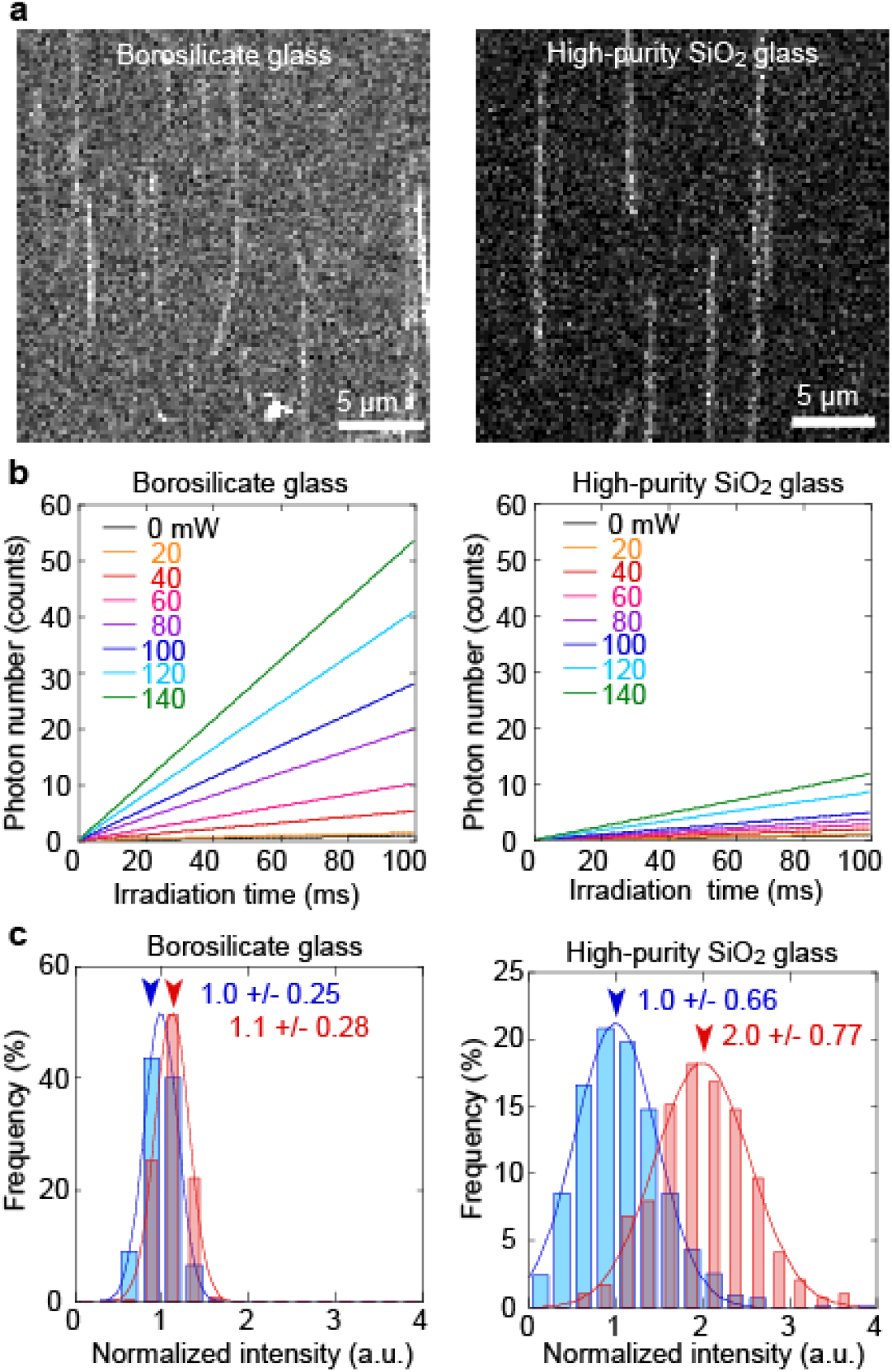
Improvement of signal-to-noise ratio in SHG measurement. (**a**) SHG images of single MTs on borosilicate glass (*left*) or on high-purity SiO_2_ glass (*right*). Scale bars are 5 μm. (**b**) Exposure time dependence of photon numbers of SHG emitted from borosilicate glass (*left*) or on high-purity SiO_2_ glass (*right*) at various source incident powers (0–140 mW). Colours indicate the powers. (c) Histograms of normalized intensity of the scattered SHG from substrate (*blue*) and MTs (*red*) using borosilicate glass (*left*) or high-purity SiO_2_ glass (*right*). The intensities were normalized to the averaged background intensity. The lines were fit to a Gaussian distribution. The values indicate (mean) +/-(standard deviation), and each mean value is indicated by an arrowhead.

### Structural differences in a single MT detected by polarization SHG microscopy

We first investigated the possibility that the electric polarization of SHG light varies according to the structure of a single MT. The α- and β-tubulin in a MT form almost indistinguishable structures, and a one-dimensional array (protofilament) of tubulin subunits with an approximately 4-nm pitch forms a parallel bundle to make a tubular structure that is approximately 25 nm in diameter. The authors of recent cryo-EM studies have reported the conformational changes in the MT^25,26^. The overall conformation of tubulin subunits is conserved, but the tilt angle of each tubulin subunit changes depending on the binding nucleotide. The tubulin subunits tilt down toward the outside and away from the tube axis when bound to a guanosine diphosphate (GDP) molecule stabilized with the anti-cancer drug paclitaxel (Taxol), compared with when they are bound to the guanosine triphosphate (GTP) analogue guanosine-5’-[(α,β)-methyleno]triphosphate (GMPCPP) (Fig. 3a). In the present study, we assumed the variation in the orientation of the tubulin subunits could be represented by a single angle parameter: the tilt angle of the permanent dipole of the tubulin subunits (*φ*). It can be assumed that the dominant hyper-polarizability lies in the same direction as the dipole. The validity of this assumption is discussed in detail in the Discussion section.

**Figure 3.**
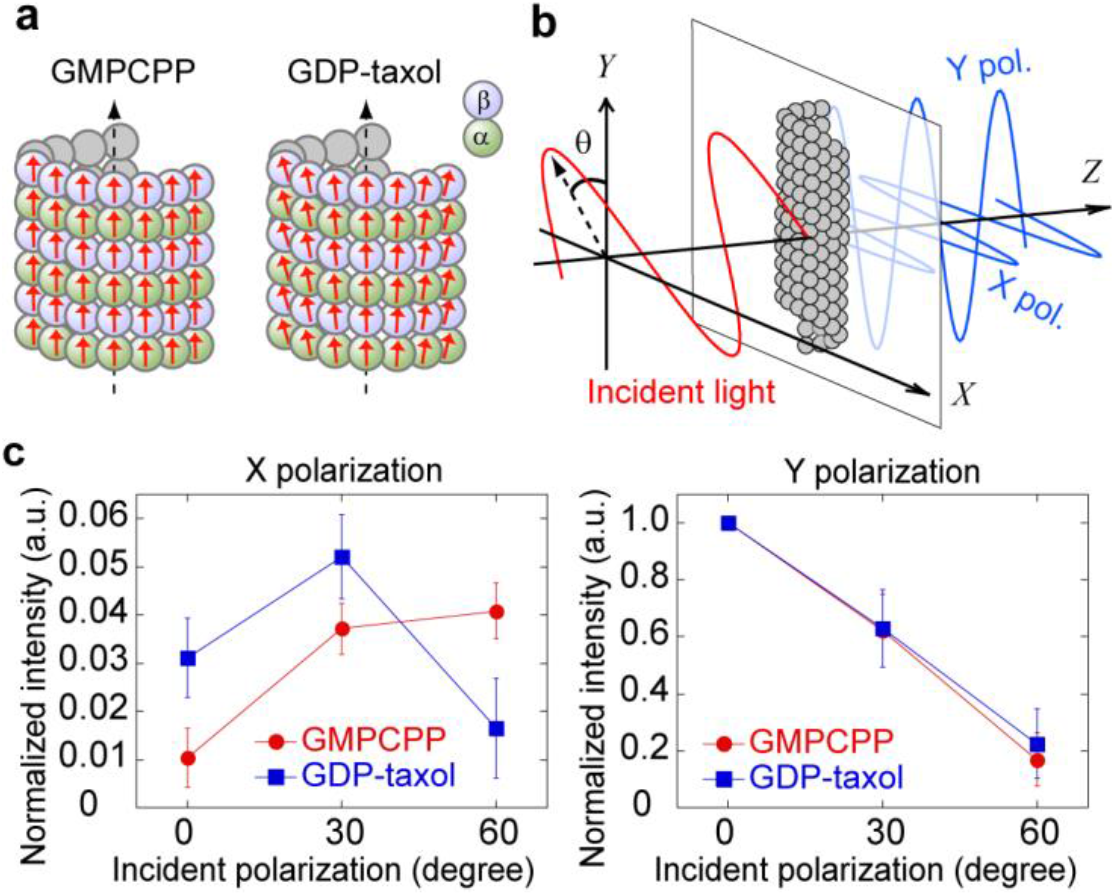
Structural analysis of a single MT by polarization SHG microscopy. (**a**) Schematic of structural differences between GMPCPP-MT and GDP-Taxol-MT. Red arrows indicate dipoles. (**b**) Orientation of an MT and incident polarization defined in the *XYZ* laboratory coordinate system. (**c**) Incident polarization dependence of SHG intensity of single MTs. *Red*, GMPCPP-MT (*N* = 255); *Blue*, GDP-MT (*N* = 100). Error bar, standard error. (**d**) SHG images of a MT bundle at incident polarizations in the range 0°–180° as indicated by arrows.

To investigate the conformational differences between GMPCPP-MT and GDP-Taxol-MT using SHG polarization, we adsorbed single fibres of GMPCPP-MT and GDP-Taxol-MT onto a glass coverslip, which we oriented along the laboratory axis *Y* (the vertical axis in the image plane), and measured the dependence of the *X*- and *Y*-polarization components of the SHG signal from the MTs on the incident polarization angle (*θ*)(Fig. 3b). There was an obvious significant difference between the two MT states in the *X*-polarization component but not in the *Y*-component (Fig. 3c), indicating that SHG microscopy is capable of sensing structural differences in an MT. It should be noted that the intensity of the *X*-polarized SHG at *θ* = 0° and *θ* = 90° should theoretically be zero under paraxial approximation, whereas it was positive in our experiment. This discrepancy can be accounted for by the depolarization effect of an objective lens with a high numerical aperture whose surface curvature causes rotation of polarization through refraction as well as propagation vector^28^ (Fig. S3, and see the Methods section). It was hypothesized that the present method proactively used the depolarization effect of the objective lens in the real system to highlight the differences.

### Theoretical calculation of the depolarization effect in the detection of the structural differences in a single MT

To evaluate the influence of depolarization on the MT polarization measurements, we calculated incident polarization dependence curves (0°–90°) with and without consideration of the depolarization effect (Fig. 4ab). A virtual MT was placed at the centre of the image plane and oriented along the *Y* axis, and the tilt angle of the dipole of the tubulin subunits (*φ*) was varied from 0° to 40° with respect to the MT axis. In the absence of the depolarization effect, it is not possible to detect the structural difference from the polarization curve of the *X*-polarized component, because the curve shape of the X-polarized component does not change with *φ* (Fig. 4a). In the presence of depolarization, the curve shape changes significantly depending on the value of *φ* (Fig. 4b). The normalized curves highlighted the structural difference between the two MT states owing to the presence of the depolarization effect (Fig. 4c). It should be noted that there is no guarantee the calculation presented here agrees with the experimental results owing to the complexity of the depolarization property of the actual objective lens.

**Figure 4.**
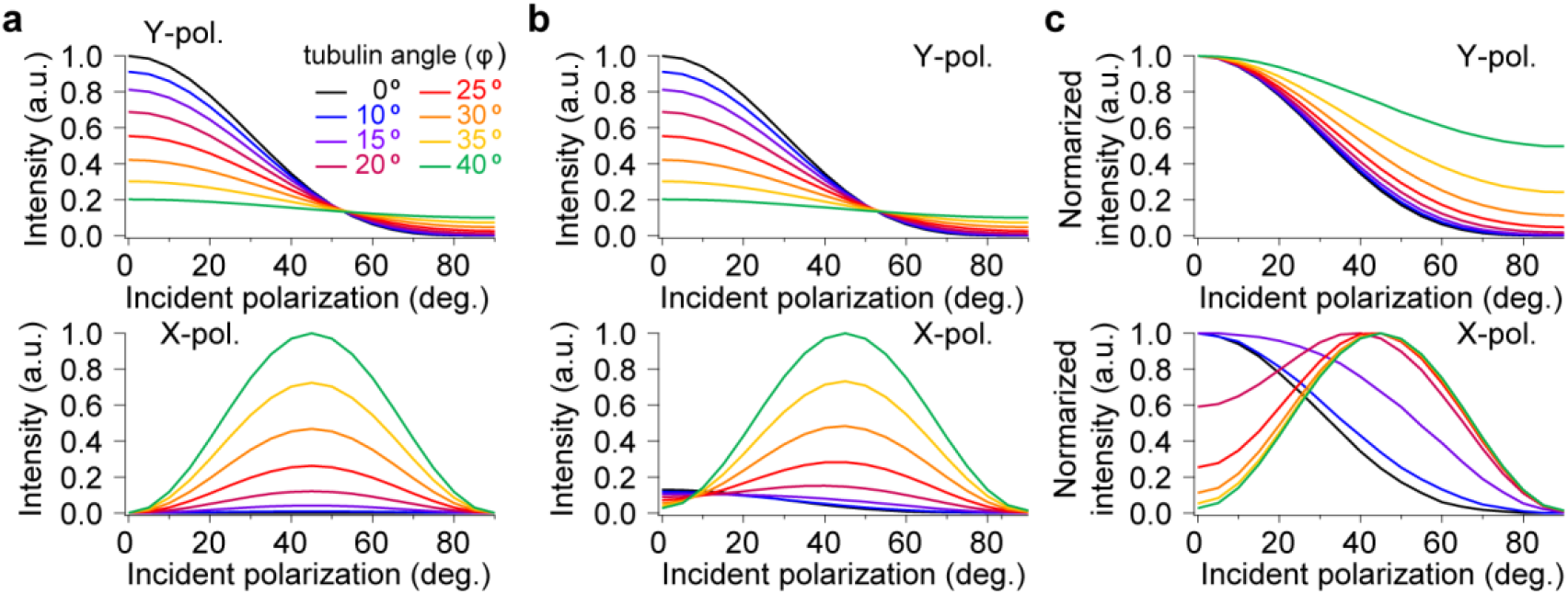
Theoretical calculation of the depolarization effect on the incident polarization dependence of SHG intensity. (**a-b**) Incident polarization dependence of *X*- and *Y*-polarized components of SHG intensity at the detector, calculated in (**a**) presence and (**b**) absence of the depolarization effect. The tubulin orientation angle *φ* varied from 0° to 40°. (**c**) Normalized curves of (**b**).

### Estimation of dipole angle of a tubulin subunit from the incident polarization dependence of SHG

The dipole angle of each tubulin subunit was calculated from the incident polarization dependence of SHG intensity. For precise measurement of the polarization dependence, we used dense bundles of parallel microtubules (MT bundles) to increase the SHG signal, which is proportional to the square of the number of tubulin subunits in a focal volume. Hundreds of microtubules were grown in a *Xenopus* egg extract from the sperm head, which served as the artificial microtubule organizing centre^29^ (Fig S4, and see the Methods section). Because the GMPCPP-MTs tended to form short filaments, and it was difficult to form large MT bundles, we used GDP-bound MT (GDP-MT) instead of GMPCPP-MT, and compared it with GDP-Taxol-MT. The angle of permanent dipole of a tubulin subunit (*φ*), as well as that of the dominant hyper-polarizability, can be derived from only two independent components of the SHG tensor, as previously described^30^, which are measurable by the incident polarization dependence of SHG intensity (Fig. 5ab). The histograms of the estimated *φ* with respect to the tube axis (Fig. 5c) suggested that Taxol changes the *φ* values of GDP-bound MT (without Taxol, 38.5°; with Taxol, 39.7°) (Fig. 5d). This tilt angle change is consistent with the structures obtained using cryo-EM^26^.

**Figure 5.**
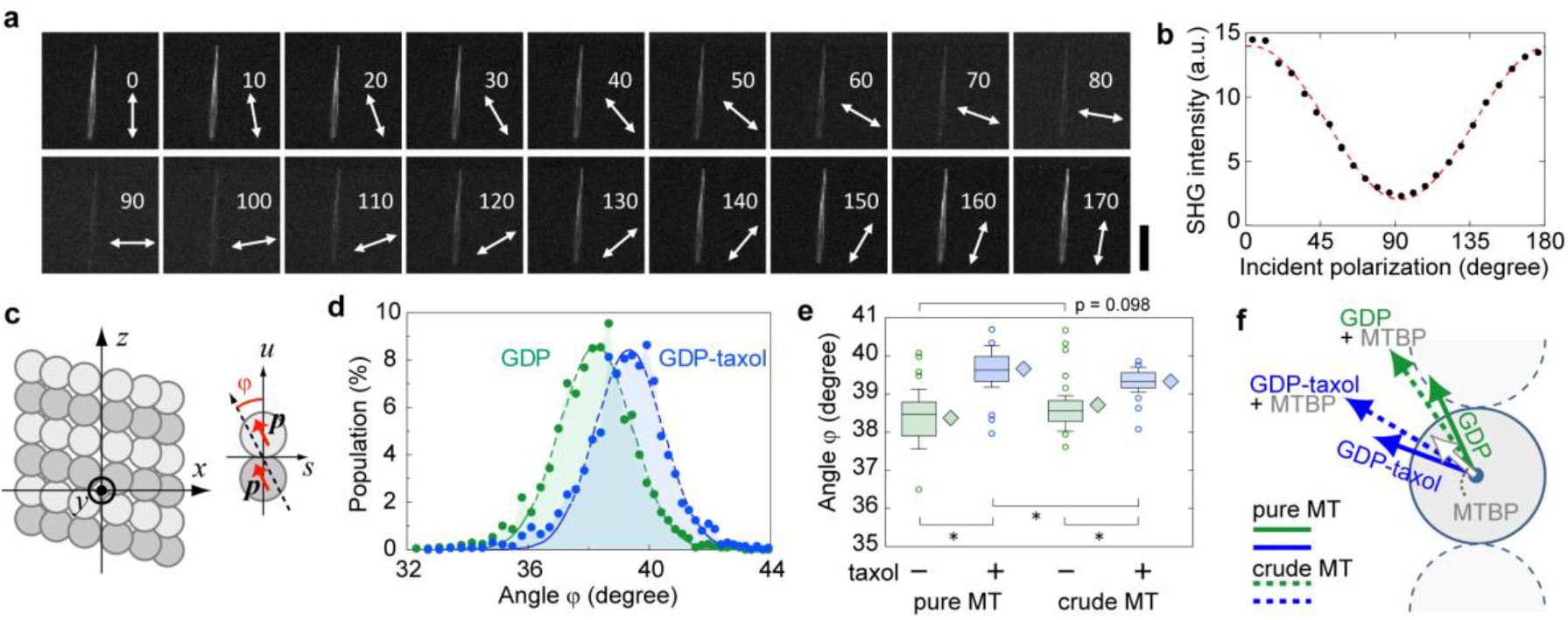
Structural analysis in MT bundle by polarization SHG microscopy. (**a**) SHG images of an MT bundle at incident polarizations in the range 0°–180° as indicated by the arrows. (**b**) A typical polarization dependence of SHG intensity of an MT bundle. Red line, theoretical curve. (**c**) Definition of axes and angle *φ*. The intra-tube axes *z* and *x* (*y*) are parallel and perpendicular to the MT axis, the intra-tubulin axes s and *u* are radial from and parallel to *z*, and *φ* is the tilt angle of tubulin subunits, ***p***, from the *z* axis (See also Fig. S7). (**d**) Histograms of estimated *φ* for GDP-MT (*green*, *N* = 2095 in 24 bundles) and GDP-Taxol-MT (*blue*, *N* = 1378 in 29 bundles) composed of pure tubulins. (**e**) Boxplots of the average *φ* in GDP-MT (*green*) and GDP-Taxol-MT (*blue*) bundles composed of pure (*left*) and crudely purified tubulins (*right*). *N* = 24, 29, 43, and 46 bundles. Asterisks indicate *p* < 0.01 (Student’s *t*-test). (**i**) Interpretation of measured dipole angles shown in (**f**). Solid and dotted arrows (*green*: GDP-MT, *blue*: GDP-Taxol-MT) indicate the dipoles of MTs composed of pure tubulins and crudely purified tubulins, respectively. The white arrow indicates the estimated dipole for MTBP.

For the MTs composed of crudely purified tubulins decorated with MT-binding proteins (MTBPs), such as microtubule-associated proteins, the estimated dipole angles were slightly larger and smaller for GDP-MT and GDP-Taxol-MT, respectively (*φ* = 38.7° for GDP-MT and 39.3° for GDP-Taxol-MT) (Fig. 5e), compared with those of the finely purified tubulins. We interpreted this small shift to mean that the average dipole angle of the MTBP is between the average dipole angles of the two states of tubulin, and accordingly shifted the estimates of the effective angle towards the dipole of the MTBP (Fig. 5f). The phenomenological interpretation of these data is included in the Discussion section.

### Temporal structural analysis in a protein crystal of photo-switchable fluorescent proteins

We next examined the feasibility of a time-lapse observation of structural changes in a protein crystal to provide a tool that complements crystallography. We chose photo-induced conformational change for a model of the dynamic phenomenon, because the timing of the conformational changes can be precisely controlled by light irradiation. For the present experiment, we chose Kohinoor, a positively photo-switchable fluorescent protein whose chromophore is activated to a fluorescent state (active state) by 470-nm irradiation, and inactivated to a non-fluorescent state (inactive state) by 395-nm irradiation^31^. We crystallized Kohinoor, because an ordered array of protein molecules effectively increases the SHG intensity, produces a high S/N ratio, and simplifies data analysis (Fig. 6ab). Both the intensity of the SHG signal and its polarization dependence differed markedly between the active and inactive states when recorded in repeated cycles of activation and inactivation (Fig. 6c, *circles*, and S5ab). The observed SHG signal also contained two-photon excited fluorescence. The polarization dependence of the fluorescence was quite different from that of SHG, and there was no difference between the two states (Fig. 6c, *rectangles*, and S5c). This indicates that the difference in the observed polarization dependence originated not from fluorescence but from SHG, owing to the structural differences in a Kohinoor crystal. The present method enabled tracking of the dynamic change in both the activation (Fig. 6d) and inactivation (Fig. 6e) processes.

**Figure 6.**
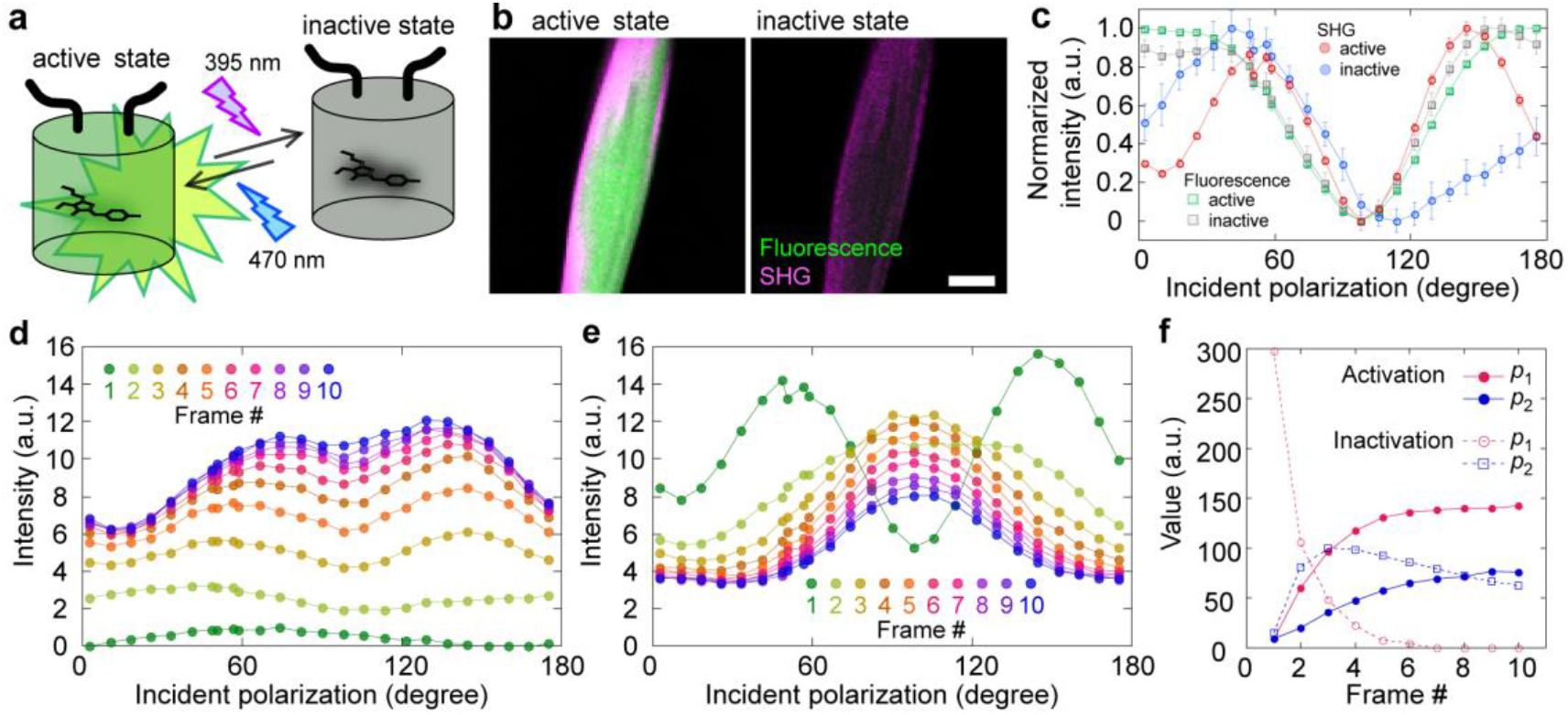
Temporal structural analysis in a protein crystal of photo-switchable fluorescent proteins. (**a**) Schematic of a state transition in photo-switchable protein, Kohinoor. (**b**) SHG (*magenta*) and fluorescence (*green*) images of a Kohinoor crystal in the active state (*left*) and the inactive state (*right*). The scale bar is 20 μm. (**c**) Incident polarization dependence of SHG of the active state (*red circles*) and the inactive state (*blue circles*) and fluorescence of the active state (*green rectangles*) and the inactive state (*grey rectangles*). The intensities were normalized from 0.0 to 1.0 by the minimum and the maximum of the average of 10 trials. (**d,e**) Time developments of the incident polarization dependence curves during 470-nm irradiation (d) and 395-nm irradiation (**e**). Colours from green to blue indicate the frame number from 0 to 10. (**f**) Time courses of results fitted to the phenomenological model (Equation S4, see Methods for details). *Red closed circles, p*_1_ in activating process; *Blue closed circles, p*_2_ in activating process; *Red open circles, p*_1_ in inactivating process; *Blue open circles, p*_2_ in inactivating process.

The crystal structure of Kohinoor has not been resolved. Therefore, we constructed a phenomenological model based on the crystal lattice structure of Padron, which is a previously reported photo-switchable fluorescent protein^32^. According to the model, two positive real parameters—*p*_1_ and *p*_2_—are defined as beneficial variants to describe the pattern of polarization dependence of SHG in the Kohinoor crystal (see the Methods section for details). The state transition of a Kohinoor crystal has been successfully represented using these parameters estimated through SHG tensor analysis (Fig. 6f and S5).

## DISCUSSION

In the present study, we improved the S/N ratio in SHG microscopy to detect the SHG signal emitted from a single MT (Fig. 1 and 2), and established a method of analysing the structural differences between MTs using SHG tensor components (Fig. 3 and 4). When the signal was sufficiently high, we were able to calculate the dipole angle of each tubulin subunit from the incident polarization dependence of SHG intensity (Fig. 5). The SHG microscope also enabled us to investigate the structural change in a crystal of a photo-switchable fluorescent protein (Fig. 6). Therefore, the SHG tensor is undoubtable coupled to the protein structure, and SHG polarization microscopy is potentially useful for the analysis of protein structures.

The background noise in the SHG measurements originated from the glass coverslips used for the specimens (Fig. 2). Although the mean value and the variance of noise intensity decreased significantly when the borosilicate substrate material was replaced with high-purity SiO_2_, the noise level persisted. This unexpected noise has not been considered in previous SHG studies of bulk samples^3,5-13,30^. The problem might be negligible when the laser focus is significantly far from the substrate, or when the signal intensity generated by the specimen is sufficiently high. We think that the generation of a periodically inverted electrostatic field via the coherent photogalvanic effect is one of the major origins of the SHG light emitted by glass^33,34^.

Fortunately, the depolarization effect of the objective lens is thought to enhance the dependence on polarization of the MT structure (Fig. 3 and 4). In other words, the depolarization effect is necessary for structural analysis using SHG microscopy. Designing optics solutions by considering depolarization over optical pathways makes SHG structural analysis more quantitative. This, together with background noise, are the next issues to be solved.

A discussion of the validity of the present estimation of the dipole angle (Fig. 5) is required. In the present study, the local coordinate system (*s,t,u*) was independently defined in each protofilament from the principal system (*x,y,z*) (Fig. S7a). The local axes s and *t* were set to radial and azimuthal directions at the location of the corresponding protofilaments, and the *u* axis was parallel to the MT axis (*u//z*). In general, the three-dimensional orientation of a dipole ***p*** in a tubulin subunit can be expressed by two angle parameters: *φ* and *δ* (Fig. S7b). The former parameter (*φ*) is, as previously mentioned, defined as the angle between *u* and ***p***, whereas the latter (*δ*) is the orientation of ***p*** within the *st* plane (= *xy* plane). Because an MT has rotational symmetry about the *z* axis, all protofilaments have the same axial dipole component (*p_A_*). Therefore, it is only necessary to discuss the lateral component *p_L_* with regard to the SHG tensor for the entire microtubule. Figure S7c illustrates nine examples of possible alignments of *p_L_* projected in the *xy* plane. The SHG tensor for an MT is calculated as the summation of the local SHG tensors for the protofilaments, and the data regarding the lateral orientation vanishes as a result of integration, even if the configuration of the dipole array varies as a function of *δ*. Therefore, in principle it is possible to obtain only a *φ* value.

The presence of MTBPs significantly affects the estimation of the angle parameter *φ* in both the GDP and GDP-Taxol states of MTs (Fig. 5e). Assuming that: (i) MTBPs on average exhibit a permanent dipole; and (ii) the permanent dipoles of tubulin subunits and MTBPs are independent of each other, the total SHG tensor (*χ*_total_) is expressed as sum of the SHG tensors of MT (*χ*_MT_) and of MTBPs (*χ*_MTBPs_). MTBPs are uniformly distributed over the entire surface of the MT with rotational symmetry (Fig. S7c). As discussed above, the contribution of *χ*_MTBPs_ is thought to originate from an average dipole of MTBP with some specific angle *φ*. Because the lateral orientations of the dipoles make no contribution to the resultant SHG tensor components, it is possible to visualise a simplified situation in which all the dipoles exist in the same plane (the *su* plane in Fig. S7b; Fig. 5f).

According to the experimental results (Fig. 5e), the orientation of the MTBP dipole (Fig. 5f, *white*) must exist in the range between those of GDP-MT (Fig. 5f, *green*) and GDP-Taxol MT (Fig. 5f, *blue*). Because the structural difference between GDP-MT and GDP-Taxol-MT is minimal, the dipoles of the two structures can be considered equivalent in magnitude. Hence, we determined the angle of the MTBP dipole to be 38.58° and the magnitude to be 0.156-fold that of the MTs. Thus, the present method is also capable of detecting the attachment/detachment of proteins on a protein filament.

We represented the structural change in a Kohinoor crystal with two parameters based on the phenomenological model (Fig. 6f). Considering the common mechanism displayed by Dronpa-based photo-switchable fluorescent proteins^35,36^—in which the benzene ring in the chromophore is dynamically flipped through a *cis–trans* transition—the same phenomenon might contribute to the change in the polarization dependence of SHG. However, an understanding of the relationship between these parameters and the structural change of the chromophore in Kohinoor require more information about the crystal structure of Kohinoor.

In the present study, we proved that SHG polarization is a label-free optical modality that can provide structural information about proteins. SHG measurements do not require labelling, unlike NMR and FRET, and are therefore applicable to long-term sequential sampling. The present study represents an important step toward the creation of a new structural analysis platform to support next-generation structural biology.

## METHODS

### Construction of the SHG microscope

Polarization-resolved SHG measurements were performed using a purpose-built laser-scanning SHG microscope in the transmission geometry with a polarization-controlled device in the incident light pathway (Fig 1). We used a mode-locked Ti:sapphire laser (Chameleon Vision II, Coherent, Inc., Santa Clara, California, USA) with a wavelength of 810 nm, a repetition rate of 80 MHz, and a pulse duration of 200 fs as an excitation light source. The laser beam was introduced to a Glan Laser polarizing prism (GL10-B, Thorlabs, Inc., Newton, New Jersey, USA) and a high-speed polarization controller composed of a pair of electro-optic modulators (Model 350-160, Model 350-80, Conoptics, Inc., Danbury, Connecticut, USA)^27^. The beam was deflected by an *x-y* scanner consisting of a pair of Galvano mirrors (VM500+, GSI, Inc., Bedford, Massachusetts, USA), and was focused by a dry objective lens with a numerical aperture (NA) of 0.95 and a magnification of 40 (CFI Plan Apo Lambda, Nikon, Inc., Tokyo, Japan). The SHG signal emitted in the forward direction was collected using another oil-immersion objective lens with an NA of 1.45 and a magnification of 100 (CFI Apo, Nikon, Inc.), and detected using a photon-counting photomultiplier tube (PMT) module (H10680-210, Hamamatsu Photonics, Inc., Shizuoka, Japan) after passing through a short-pass filter (FF01-680/SP, Semrock, Inc., Rochester, New York, USA) and a bandpass filter (FF01-405/10-25, Semrock, Inc.). For the investigation of single MTs, the SHG signal was split into *X-* and *Y*-polarized components by a polarizing beam-splitter, and detected by two PMT modules. However, the beam splitter was not used to investigate MT bundles. Therefore, the total SHG intensity was detected using a single PMT module. To demonstrate imaging with our SHG microscope, we examined nerve cells prepared by fixing microtubules with Taxol followed by extraction of other proteins and organelles using Triton X-100. As shown in Fig. S1, bundled MTs and the polarization of a small centrosome were selectively examined in dendrites and axons without labelling.

### Fluorescent staining of tubulin

Tubulin was purified from a porcine brain through four cycles of polymerization and depolymerization using 1 M PIPES buffer (1 M PIPES-KOH, 1 mM EGTA, 1 mM MgCl_2_, pH 6.8) for the effective removal of microtubule-associated proteins^37^. The labelled tubulin was prepared by incubating the polymerized microtubule with Alexa Fluor 647 NHS Ester (A-20106, Life Technologies, Inc.) for 30 min at 37°C^38^. We then purified the functionally labelled tubulin with two cycles of polymerization and depolymerization.

### Preparation of single microtubules for observation

Fluorescently labelled tubulin was mixed with unlabelled tubulin to yield a final labelling stoichiometry of 2%. For the GMPCPP microtubules, 2.0 μM tubulin was incubated in PEM (100 mM Pipes-KOH, 1 mM ethylene glycol-bis(2-aminoethylether)-N,N,N’,N’-tetraacetic acid (EGTA), 1 mM MgCl_2_, pH 6.9) with 0.2 mM GMPCPP and 0.2 mM MgSO_4_ at 37°C for 30 min. For the GDP-Taxol microtubules, 30 μM tubulin was first polymerized by incubation in PEM with 1 mM GTP and 1 mM MgCl_2_ at 37°C for 15 min, and the polymerized microtubules were then resuspended in PEM with 0.5 mM GDP, 0.5 mM MgSO_4_, and 10 μM paclitaxel. A rectangular flow channel between two glass coverslips (borosilicate or high-purity SiO_2_) had a typical width of 2 mm, a height of 5 μm, and a length of 18 mm. The surface was coated with 25 mg/mL poly-L-lysine dissolved in 100 mM borate buffer (pH 8.5) to enhance the electrostatic attachment of the negatively charged microtubules to the glass surface.

### Measurement and analysis of polarization dependence of single microtubules

Polarization-resolved SHG imaging was carried out several times for each of the GMPCPP-MT and GDP-Taxol-MT samples. The laser focus was raster-scanned in a 25 × 25 μm^2^ square with a pixel pitch of 250 nm (100 × 100 pixels) and a dwell time of 30 ms. The incident polarization was swept within the dwell time at every pixel position. The average laser power at the focal plane was approximately 30 mW, and the corresponding peak power density of a pulse at the focus was estimated to be 200 GW/cm^2^. We chose straight microtubules with unique orientation along the *Y*-axis from the SHG images. Photon-counting data in the area covering each microtubule were integrated so that we ultimately obtained one set of polarization-dependent data per element. We also extracted the integrated photon-count in a blank region to subtract the background noise. After the background noise had been subtracted, we obtained SHG intensities *I_X_*(*θ*) and *I_Y_*(*θ*) from one dataset, in which *I_X_, I_Y_*, and *θ* were the *X*-polarized intensity, the *Y*-polarized intensity, and the incident polarization angle, respectively. For analysis, we chose 0°, 30°, and 60° as the values of *θ*. Finally, the polarization-dependent SHG intensities were linearly normalized so that *I_Y_*(0°) equalled 1.0.

### Theoretical calculation of the depolarization effect of the collection objective lens

We calculated the direction of the radiation from an *X*-polarized dipole collected using an objective based on electrodynamics theory^39^. The objective was assumed to be an ideal lens with high numerical aperture, which collimated a spherical wave diverging from its focal position. The refraction of vectorial rays was calculated at all positions on the lens plane based on Snell’s law^28^, and the rays were projected on the Fourier plane (Fig. S3a). To simplify calculation, an aberration-free singlet lens with a maximum detection angle of 60° was used instead of an objective lens. An SHG dipole with a wavelength of 405 nm was placed at the geometrical focus. The detector plane was divided into small pixels, and the polarization state of the optical rays from the dipole was calculated in all the pixels.

Looking at the intensity distribution of the *X*- and *Y*-polarized components on the Fourier plane, although the light source was purely *X*-polarized, *Y*-polarized light was detected particularly at the outer region (high-NA region) of the lens (Fig. S3b, *left*). Depolarization was more obvious at the periphery of the Fourier plane. More precisely, the *Y*-polarized component was zero on both the *X*-axis (*Y* = 0) and *Y*-axis (*X* = 0) because the rays on the axes were purely *p*-polarized (on the *X*-axis) or *s*-polarized (on the *Y*-axis). In contrast, the *Y*-polarized component intensity became large at the diagonal axes *Y* = ± *X*. This is clearly visualized by the arrows on the plane indicating the directions of polarization (Fig. S3b, *right*).

To evaluate the influence of depolarization on the measurement of MT polarization (Fig. 4), we assumed virtual MTs with various *φ* values (tilt angle of tubulin subunits) were placed on the focal plane with orientation along the *Y*-axis. The induced polarization of SHG was calculated using the SHG susceptibility tensor, as described below. The intensity of the *X*- and *Y*-polarized components was integrated at the Fourier plane to draw the polarization curves.

### Preparation of microtubule bundles for observation

*Xenopus* egg extract and demembrenated sperm heads were prepared as reported previously for the investigation of MT bundles^40^. The reactions were conducted at 37°C in rectangular flow channels with a typical width of 1.2 mm, a height of 30 μm, and a length of 18 mm, formed between two coverslips according to the following procedure (Fig. S4). The demembranated *Xenopus* sperm was incubated for 5 min in the flow channel. The floating sperm was washed with 5 μL of 7 mg/mL casein dissolved in 50 mM Pipes-KOH, 2 mM MgSO_4_, and 1 mM EGTA; pH 6.7. *Xenopus* egg extract (5 μL) was added and the mixture was incubated for 10 min to create microtubule organization centres. Unnecessary egg extract was removed by washing with 5 μL of warmed high-concentration Pipes buffer (500 mM Pipes-KOH, 10 mM MgCl_2_, 5 mM EGTA, pH 6.94) and 10 μL of warmed PEM with 30 % (v/v) glycerol (PEM-glycerol). Tubulin (8 μL, 15 μM) purified from porcine brain was loaded and incubated for 20 min in PEM with 1 mM GTP and 1 mg/mL casein. The polymerized microtubules were bundled by the flow of 20 μL of PEM-glycerol. For the Taxol-bound MTs, the washing buffer was replaced with 20 μL of PEM-glycerol with 50 μM paclitaxel.

### Polarization-resolved SHG imaging of microtubule bundles

A set of polarization-resolved SHG images was obtained by synchronous control of the *x-y* scanner and the polarization controller. The laser focus was raster-scanned in a 50 × 50 μm^2^ square with a pixel pitch of 500 nm (100 × 100 pixels) and a dwell time of 13 ms. The incident polarization was swept within the dwell time at every pixel position. The angle pitch was 7.5° and the number of angles was 26, which covered a total angle range of over 180°. The total time to obtain a whole image set was 130 s. The average laser power at the focal plane was approximately 30 mW.

### Estimation of dipole angle of a tubulin subunit from the incident polarization dependence of SHG

The SHG intensity in an area that covered an entire bundle at each polarization-resolved image was integrated to obtain one set of polarization-dependent data for one bundle. The number of integrated pixels for a set of data was in the range 100–2000 depending on the size of the bundle. The polarization-dependent data were then fitted using the following theoretical function of incident polarization angle *θ* with three fitting parameters *α*, *χ_zzz_* and *χ_zxx_*.

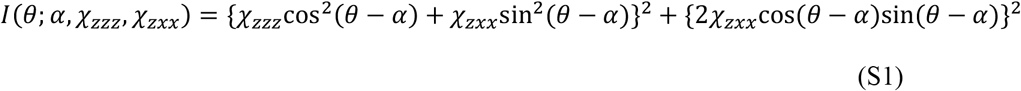

*α* denotes the average orientation angle of an MT bundle. Angles *θ* and *α* were defined according to the *Y* laboratory axis (the vertical axis in the SHG images in Fig. 3b). Parameters *χ_zzz_* and *χ_zxx_* are two components of the SHG susceptibility tensor. Equation S1 is a well-established form of SHG polarization dependence in biological filaments^30^.

We herein describe how to estimate the effective tilt angle (*φ*) of tubulin subunits (α- and β-tubulins), defined in Fig. S7, from the parameters *χ_zzz_* and *χ_zxx_*. The coordinate system for the definition of *χ_zzz_* and *χ_zxx_* is shown in Fig. 5c in the main text. The *z*-axis is defined as the direction parallel to the central axis of the MT, and the *x*- and *y*-axes are defined as perpendicular to the MT axis. Assuming cylindrically symmetric alignment of the tubulin, *χ_zzz_* and *χ_zxx_* are the only two independent non-zero components that express all other non-zero components. We applied Kleinman’s law^1^, which was appropriate because the wavelengths used in our measurements were far from the absorption wavelength of proteins in an MT. The two components are related to *φ* by the following equations:

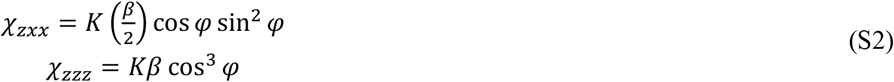

*β* denotes the magnitude of the dominant hyper-polarizability of a tubulin subunit, and *K* is a constant that is proportional to the net molecular number in the focal volume. The hyper-polarizability, *β*, is assumed to have a direction identical to the electric dipole of a tubulin subunit. The effective tilt angle (*φ*) has the following relationship with the ratio of *χ_zzz_* and *χ_zxx_*:

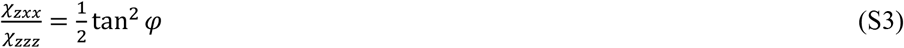

We estimated the effective tilt angle *φ* throughout this study based on the values of *χ_zzz_* and *χ_zxx_* extracted from the curve fitting Eq. (S1) to the experimental results.

### Crystallization of photo-switchable fluorescent protein Kohinoor

The cDNAs for Kohinoor kindly provided by Prof. T. Nagai were inserted into an *Escherichia coli* expression vector pAL7 (BIO-RAD, CA) and transformed into the *E. coli* strain Rosseta2 (DE3) (Merck Millipore, DE). The expressed Kohinoor was purified using a Profinity eXactTM Fusion-Tag system (BIO-RAD, CA) following the protocol recommended by BIO-RAD. The yield of purified Kohinoor was 3.0–4.0 mg. Finally, the Kohinoor solution was concentrated to 8 mg/mL in 1 mM 4-(2-hydroxyethyl)-1-piperazineethanesulfonic acid (HEPES) (pH 7.5) using Amicon Ultra Centrifugal Filters (Merck Millipore, DE).

The purified Kohinoor was crystallized using the hanging-drop vapor-diffusion technique. The Kohinoor proteins were precipitated and crystalized within 1 day at 20°C from drops prepared by mixing 1.2 μL of the protein solution with 1.2 μL of a reservoir solution comprising 50 mM HEPES (pH 8.0), 10% PEG4000, and 26.7 mM MgCl_2_.

### SHG observation of Kohinoor crystal

We introduced a 395/470-nm LED light source (Light Engine Spectra, Lumencor, Inc., Beaverton, Oregon, USA) to activate and inactivate Kohinoor, and a bandpass filter (FF01-525/45-25, Semrock) for fluorescence observation, into the present SHG microscope. Small pieces of Kohinoor crystal were loaded into a glass chamber with the buffer (phosphate-buffered saline). The SHG signal of the Kohinoor crystal was recorded using the same procedure described in the section describing the polarization dependence measurement of the microtubule bundles, except that the dwell time was 1.0 ms and the number of polarization angles was 25.

For comparison of the polarization dependence of total SHG intensity between fluorescent and non-fluorescent states (Fig. 6c), the sequence of [10 s irradiation of 470 nm light at 0.43 mW, SHG recording, 10 s irradiation of 395 nm light at 0.16 mW, SHG recording] was repeated 10 times. For time course measurements of the incident polarization dependences during 470 nm light irradiation for activation (or during 395 nm light for inactivation) (Fig. 6de), the sequence of [1.0 s irradiation of 470 nm light at 0.43 mW (1.0 s irradiation of 395 nm light at 0.016 mW), SHG recording] was repeated 10 times.

### Phenomenological model fitting the SHG observations of the Kohinoor crystal

We assumed that the space group of our Kohinoor crystal samples was *P*2_1_2_1_2 according to a report of the crystal structure of a similar positively switchable protein, Padron, into which some mutations were introduced to develop Kohinoor^31^. The SHG tensors under these symmetry conditions have only three components (*χ_xyz_*, *χ_yzx_*, and *χ_zxy_*) in the tensor principal coordinate system (*x,y,z*)^41^. We introduced an imaginary part into each component of the SHG tensor to adjust for the effect of resonance on SHG, because the SHG wavelength in the present measurement system (405 nm) was within the absorption band of Kohinoor.

The polarization dependence of the total SHG intensity was expressed as:

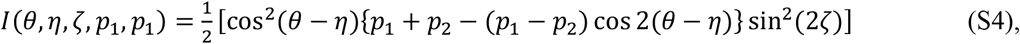

where *η* and *ζ* denote the orientations of the sample as defined in Fig. S8, and *p*_1_ and *p*_2_ are material-dependent parameters expressed with *χ_xyz_*, *χ_yzx_*, and *χ_zxy_* by the following equations:

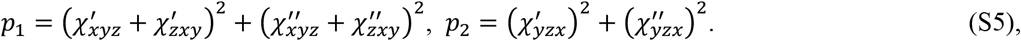

where the single and doubled primes represent real and imaginary parts, respectively.

According to equations S4 and S5, the complexity of the resonance effect was invisible under the experimental conditions when we measured total intensity. Moreover, only two real positive values of *p*_1_ and *p*_2_ were significant and beneficial to a discussion of the patterns of the polarization dependence curves of SHG from the Kohinoor crystal. Because the values of *p*_1_ and *p*_2_ can be uniquely determined as long as sin^2^(2*ζ*) is not zero, we assumed *ζ* = π/4 in the present study. In the SHG signal obtained, *I*(*θ*) comprises the polarization dependence of pure SHG intensity, *I*_SHG_(*θ*), and that of fluorescence, *I*_fl_(*θ*), as follows:

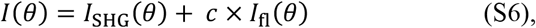

where *c* is a dimensionless parameter. The effect of fluorescence can be removed by using this equation with *I*_fl_(*θ*) detected at 525 nm. Figure S6 shows the time courses of raw data *I*(*θ*) (a,c) and *I*_SHG_(*θ*) (b,d) obtained by subtracting *I*_fl_(*θ*) from *I*(*θ*) in the activation (a,b) and inactivation processes (c,d). Hence, the parameters, *η*, *p*_1_, *p*_2_, and *c*, were estimated by least-square fitting with equations S4 and S6.

*Note: supplementary information and source data files are available in the online version of this paper*.

## ACKNOWLEDGEMENTS

We are deeply grateful to Kylius Wilkins (RIKEN, QBiC) for a critical assessment of this manuscript. We also thank Dr. A. Takai and Dr. T. Kambara (RIKEN, QBiC), and Dr. H. Inomata (RIKEN, CDB) for the preparation of the *Xenopus* egg extracts and sperm; and S. Xu, J. Asada, and M. Kakiuchi (RIKEN, QBiC) for their technical and secretarial assistance. The Kohinoor plasmid was kindly provided by Dr. D. K. Tiwari and Prof. T. Nagai (Osaka University). This work was supported by the Special Postdoctoral Researchers’ Program (SPDR) of RIKEN (to J.K. and T.S.); a Grant-in-Aid for JSPS Research Fellows (to J.K.); Grants-in-Aid for Scientific Research (KAKENHI; grant numbers 25871144 to J.K.; 16H05119, 15H01334, 26115721, and 25293046 to Y.O.; and 16H01439 to T.I.) from the Ministry of Education, Culture, Sports, Science, and Technology; the Uehara Memorial Foundation (to Y.O.); the Takeda Science Foundation (to Y.O.); the All RIKEN Research Project on Single Cells (to Y.O. and T.M.W.); and the RIKEN Pioneering Project on Dynamic Structural Biology (to Y.O.).

## AUTHOR CONTRIBUTIONS

T.M.W. and Y.O. conceived the experiments. J.K. built the microscope system and performed the experiments. T.S. and Y.O. designed the microtubule experiments. M.T. and K.I. crystalized the Kohinoor protein. T.I. performed the simulation and supervised the work. T.M.W. designed the project. J.K. and T.M.W. wrote the manuscript with discussion and feed-back from all the authors.

## COMPETING FINANCIAL INTERESTS

The authors declare no competing financial interests.

*Information about reprints and permissions is available online at http://www.nature.com/reprints/index.html*.

